# Variation in the plasma membrane monoamine transporter (PMAT, encoded in *SLC29A4*) and organic cation transporter 1 (OCT1, encoded in *SLC22A1*) and gastrointestinal intolerance to metformin in type 2 diabetes: an IMI DIRECT study

**DOI:** 10.1101/436980

**Authors:** Adem Y Dawed, Kaixin Zhou, Nienke van Leeuwen, Anubha Mahajan, Neil Robertson, Robert Koivula, Petra JM Elders, Simone P Rauh, Angus G Jones, Reinhard W Holl, Julia C Stingl, Paul W Franks, Mark I McCarthy, Leen ‘t Hart, Ewan R Pearson, for the IMI DIRECT Consortium.

## Abstract

**Objectives:** 20-30% of patients with metformin treated type 2 diabetes experience gastrointestinal side effects leading to premature discontinuation in 5-10% of the cases. Gastrointestinal intolerance may reflect localised high concentrations of metformin in the gut. We hypothesized that reduced transport of metformin into the circulation via the plasma membrane monoamine transporter (PMAT) and organic cation transporter 1 (OCT1) could increase the risk of severe GI side effects.

**Research Design and Methods:** The study included 286 severe metformin intolerant and 1128 tolerant individuals from the IMI DIRECT consortium. We assessed the association of patient characteristics, concomitant medication and the burden of mutations in the *SLC29A4* and *SLC22A1*, genes that encode PMAT and OCT1, respectively, on odds of metformin intolerance using a logistic regression model.

**Results:** Women (p < 0.001) and older people (p < 0.001) were more likely to develop metformin intolerance. Concomitant use of metformin transporter inhibiting drugs increased the odds of intolerance by more than 70% (OR = 1.72 [1.26-2.32], p < 0.001). In a logistic regression model adjusted for age, sex, weight and population substructure, the G allele at rs3889348 (*SLC29A4*) was associated with GI intolerance (OR = 1.34[1.09-1.65], p = 0.005). rs3889348 is the top cis-eQTL for *SLC29A4* in gut tissue where carriers of the G allele had reduced expression. Homozygous carriers of the G allele treated with metformin transporter inhibiting drugs had over three times higher odds of intolerance compared to carriers of no G allele and not treated with inhibiting drugs (OR = 3.23 [1.71-6.39], p < 0.001). Using a genetic risk score (GRS) derived from *SLC29A4* (rs3889348) and previously reported *SLC22A1* variants (M420del, R61C, G401S), the odds of intolerance was more than twice in individuals who carry three or more risk alleles compared with those carrying none (OR = 2.15 [1.20-4.12], p = 0.01).

**Conclusions:** These results suggest that intestinal metformin transporters and concomitant medications play an important role in gastrointestinal side effects of metformin.

## Introduction

Metformin therapy can cause gastrointestinal discomfort that negatively affects quality of life and adherence to prescribed medications. Gastrointestinal side effects usually manifest as nausea, vomiting, diarrhoea, flatulence, indigestion, bloating, abdominal discomfort and stomach ache. 20-30% of metformin treated participants with type 2 diabetes experience gastrointestinal side effects leading to premature discontinuation in 5-10% of the cases (1, 2). This inhibits adherence to therapy and may lead to a change of treatment, depriving intolerant patients of effective diabetes therapy. Despite its clinical importance, the underlying pathophysiology of metformin intolerance is not yet clear. However multiple possible hypotheses have been proposed including high intestinal metformin concentration (3, 4), its effect on the gut microbiota (5), altered transportation of serotonin or direct serotonergic effects (6), and reduced ileal absorption of bile acid salts (7).

Metformin is not metabolized and is excreted unchanged in the urine. At physiologic pH, it is hydrophilic due to the presence of a quaternary ammonium group that results in a net positive charge. Therefore, Metformin does not efficiently diffuse across the biological membranes and requires carrier-mediated transport. Multiple solute carrier transporters expressed in membranes of the enterocytes, hepatocytes and the kidney are reported to be involved in the absorption, distribution and elimination of metformin. Metformin requires the entire length of the small intestine to be absorbed (8): around 20% of the administered dose is absorbed in the duodenum and 60% in the jejunum and ileum. The remainder reaches the colon and remains unabsorbed. PMAT and OCT1 are reported to play the major role in the intestinal absorption of metformin (9). While PMAT is expressed in the apical (luminal) membrane of the enterocytes, intestinal localization of OCT1 is ambiguous (9–11). An association between reduced function alleles in *SLC22A1* and concomitant use of OCT1 inhibiting drugs with metformin intolerance has been reported (12, 13). An interaction between OCT1 and Serotonin Transporter (SERT) has also been shown to play an important role in the pathophysiology of metformin intolerance (13).

Whilst PMAT shares extensive substrate and inhibitor overlap with OCTs (14), there are no studies investigating its role in metformin intolerance. Therefore, we hypothesized that reduced transport of metformin by PMAT and/or OCT1 could increase intestinal metformin concentration and subsequently increase the risk of GI side effects. To address this, we used prescribing, biochemistry and clinical data from 286 metformin intolerant and 1128 tolerant individuals from the IMI DIRECT (DIabetes REsearCh on patient straTification) consortium (15).

## Research Design and Methods

### Study population

286 metformin intolerant (cases) and 1128 metformin tolerant (controls) subjects were identified from prescribing data in the IMI DIRECT consortium from participating centres across northern Europe (15). Each participant consented to participate in the study and ethical approval was obtained from the medical ethics committees of the respective centres.

All metformin intolerant (cases) and metformin tolerant (controls) had: type 2 diabetes diagnosed clinically, a creatinine clearance ≥ 60 mL/min at metformin exposure, and were white Europeans aged between 18-90 years at recruitment.

### Definition of metformin intolerance

The metformin intolerance phenotype was defined in two ways: firstly, individuals who switched to an alternative agent within 6 months of stopping metformin (including modified release metformin) after having had up to 1000 mg daily metformin for up to 6 weeks, who also reported gastrointestinal side effects on the metformin treatment as the reason for switching or where gastrointestinal side effects were clearly documented in the clinical record as a reason for transfer. In an alternative definition, intolerant individuals were defined as those who could not increase their metformin immediate release dose above 500 mg daily despite an HbA1c > 7% (53 mmol/mol) and who either reported gastrointestinal side effects on more than 500 mg, or where gastrointestinal side effects were clearly documented in the clinical record as a reason for transfer.

Where the patient was asked to recall side effects, the intolerant event was limited to be within the last 5 years; if side effects were documented from clinical records then there was no time limit. Participants who did not recall being on metformin or having side effects were excluded (unless clearly documented in clinical records).

### Definition of metformin tolerance

Metformin tolerant individuals were defined as those treated with ≥ 2000 mg of metformin per day for more than a year (excluding modified release formulations of metformin) and report no side effects.

### Clinical covariates

Weight, height and creatinine were defined as the closest measured values within 180 days prior to the index intolerance event (ITE) and BMI was calculated as weight in kg / (height in m)^2^. The ITE was defined as the date when patients report gastrointestinal symptoms of metformin intolerance for cases, and for controls it is the date when patients start 2000 mg of metformin. Daily dose was the last dose during ITE for cases, and it was determined as the mean dose of prescriptions encashed during the first six months of metformin therapy for controls.

### Concomitant medications

Gut metformin transporters have strong substrate and inhibitor overlap (16). Therefore, we identified medications prescribed together with metformin previously reported to inhibit the PMAT and/or OCTs, proteins that mediate transmembrane trafficking of their target molecules, and are required for metformin absorption in the gut. These drugs are selected based on their reported half-maximal inhibitory concentration (IC_50_) values. Accordingly the use of any of the following medications with metformin were investigated: tricyclic antidepressants (TCAs) (17, 18), proton pump inhibitors (PPIs) (19), citalopram (18), verapamil (17, 18), diltiazem (18), doxazosin (17, 18), spironolactone (17, 18), clopidogrel (20), rosiglitazone (21), quinine (18), tramadol (18, 22), codeine (23), dysopyramide (24), quinidine (21), repaglinide (21), propafenone (17), ketoconazole (17), morphine (22, 23), tropisetron (25), ondasetrone (25), antipsychotic agents (17) and tyrosine kinase inhibitors (26).

### Genotyping

DNA samples from participants were genotyped at the University of Oxford (UOXF) using the Illumina Human Core Exome chip v1.0 (HCE24 v1.0). Genotype calling was performed using the GenCall algorithm in the GenomeStudio software supplied by Illumina. Data were subjected to a series of standard quality control analyses in order to highlight poorly performing genetic markers and samples prior to imputation.

Samples were excluded for any of the following reasons: call rate less than 95%, heterozygosity greater than 4 standard deviations (SD) from the mean, high correlation to another sample (pi-hat ≥ 0.2) or identified as ethnic outlier from constructed axes of genetic variation from principal components analysis implemented in the Genome-wide Complex Trait Analysis (GCTA) software (v1.24.7) (27) using the 1000 Genome as a reference. Further filtration was performed to remove: non-autosomal markers, duplicate markers (sharing the same positions), markers with minor allele frequency (MAF) <1%, Hardy–Weinberg equilibrium (HWE) p-value < 0.0001 and call rate < 98%. Imputation to the 1000 Genomes Phase 3 CEU reference panel was performed with ShapeIt (v2.r790) (28) and Impute2 (v2.3.2) (29).

### Single nucleotide polymorphism selection

As there are no functionally characterised common nonsynonymous SNPs in the *SLC29A4* gene, the tagging intronic SNPs, rs3889348 and rs2685753 (r^2^ = 0.57, D’ = 1) had been previously shown to be associated with trough steady state metformin concentration (30). Therefore, the rs3889348 G>A genotype, was extracted from existing genome-wide data. The frequency of the minor allele (A) of rs3889348 was 38%. Data for previously reported missense OCT1 variants (M420del, R61C, G401S) were also extracted from the genome-wide data. There was no deviation from HWE for any polymorphism (p > 0.05). were obtained from existing genome-wide data.

### Statistical methods

Categorical data are presented as frequency (percentage) and continuous variables as mean ± SD if normally distributed or as median and inter quartile range (IQR) otherwise. Students t-test and the Mann-Whitney U test were used to compare differences in quantitative variables distributed normally or not, respectively. Comparison of categorical variables between cases and controls was done using *X*^2^ test. Logistic regression was used to estimate the association of independent variables with metformin intolerance. Multivariate logistic regression analyses of metformin intolerance were performed assuming an additive genetic model, with all the covariates included using SNPTEST (version 2.5.2) (31). A two-tailed p-value less than 0.025 was considered statistically significant.

## Results

### Phenotypic differences between tolerant and intolerant subjects

The characteristics of tolerant and intolerant subjects are presented in Table 1. Women (p < 0.001) and older people at diagnosis or at ITE (p < 0.001) were more likely to be metformin intolerant. Compared to tolerant subjects, metformin intolerant individuals had lower weight (p < 0.001), lower creatinine clearance (p = 0.036) and were treated with a lower metformin dose (p < 0.001).

**Table 1.**
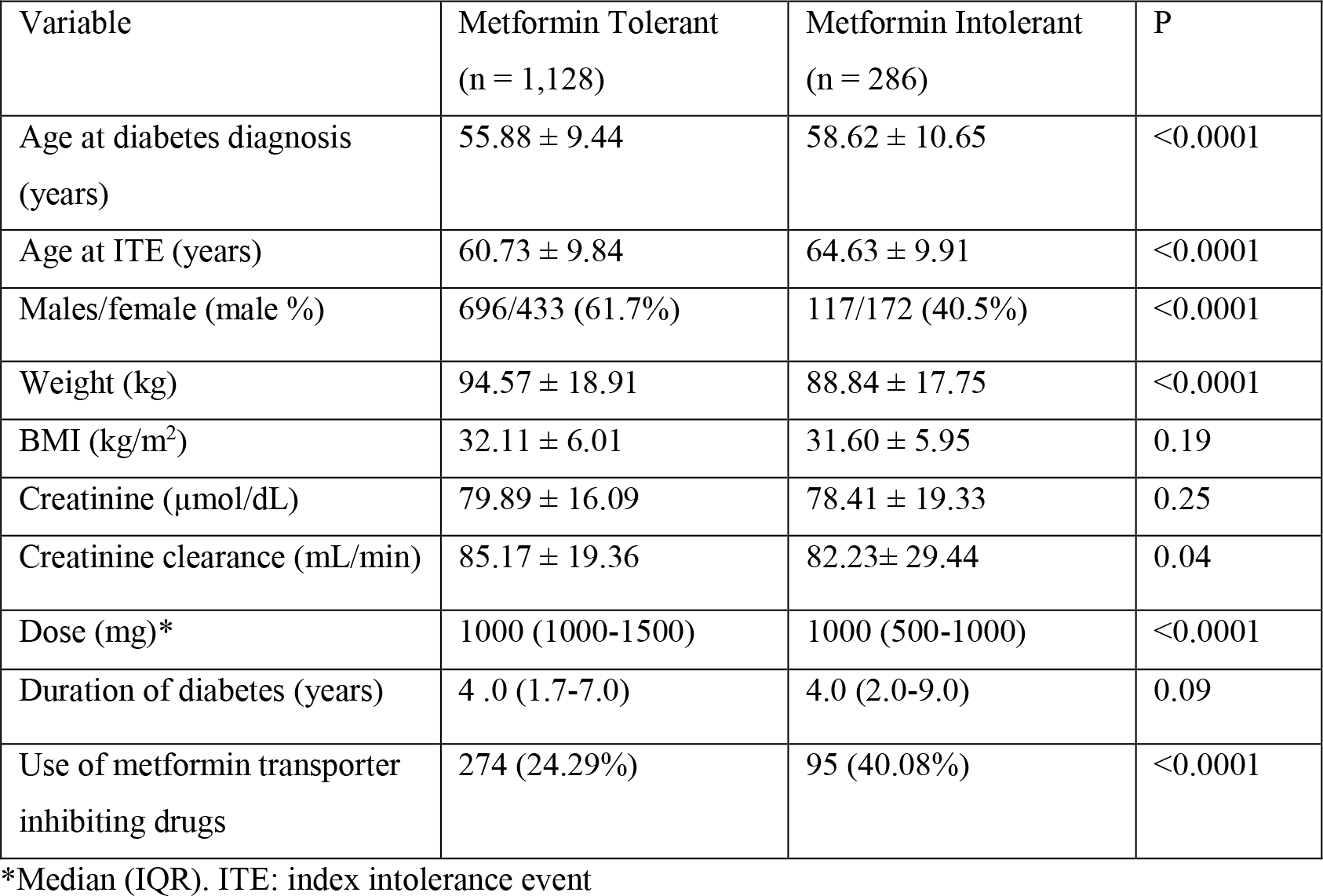
Baseline characteristics of metformin tolerant and intolerant subjects.

| Variable | Metformin Tolerant<br>(n = 1,128) | Metformin Intolerant<br>(n = 286) | P |
| --- | --- | --- | --- |
| Age at diabetes diagnosis<br>(years) | 55.88 ± 9.44 | 58.62 ± 10.65 | <0.0001 |
| Age at ITE (years) | 60.73 ± 9.84 | 64.63 ± 9.91 | <0.0001 |
| Males/female (male %) | 696/433 (61.7%) | 117/172 (40.5%) | <0.0001 |
| Weight (kg) | 94.57 ± 18.91 | 88.84 ± 17.75 | <0.0001 |
| BMI (kg/m<sup>2</sup>) | 32.11 ± 6.01 | 31.60 ± 5.95 | 0.19 |
| Creatinine (μmol/dL) | 79.89 ± 16.09 | 78.41 ± 19.33 | 0.25 |
| Creatinine clearance (mL/min) | 85.17 ± 19.36 | 82.23 ± 29.44 | 0.04 |
| Dose (mg)* | 1000 (1000-1500) | 1000 (500-1000) | <0.0001 |
| Duration of diabetes (years) | 4 .0 (1.7-7.0) | 4.0 (2.0-9.0) | 0.09 |
| Use of metformin transporter<br>inhibiting drugs | 274 (24.29%) | 95 (40.08%) | <0.0001 |

\*Median (IQR). ITE: index intolerance event

### Concomitant medications and intolerance

This analysis was performed on 233 metformin intolerant and 1128 tolerant subjects who had complete data on history of concomitant medications. Forty percent of metformin intolerant subjects were taking one or more cation transporter inhibitory drugs compared to 24% in tolerant subjects (p < 0.0001) (Table 1). In a logistic regression model adjusted for age and sex, concomitant use of these drugs increased the odds of being intolerant by 70% (OR = 1.70 [1.24-2.29], p < 0.001).

When the individual drug or drug groups were explored, concomitant use of metformin with either PPIs, TCAs or codeine increased the odds of metformin intolerance significantly (Figure1).

**Figure 1.**
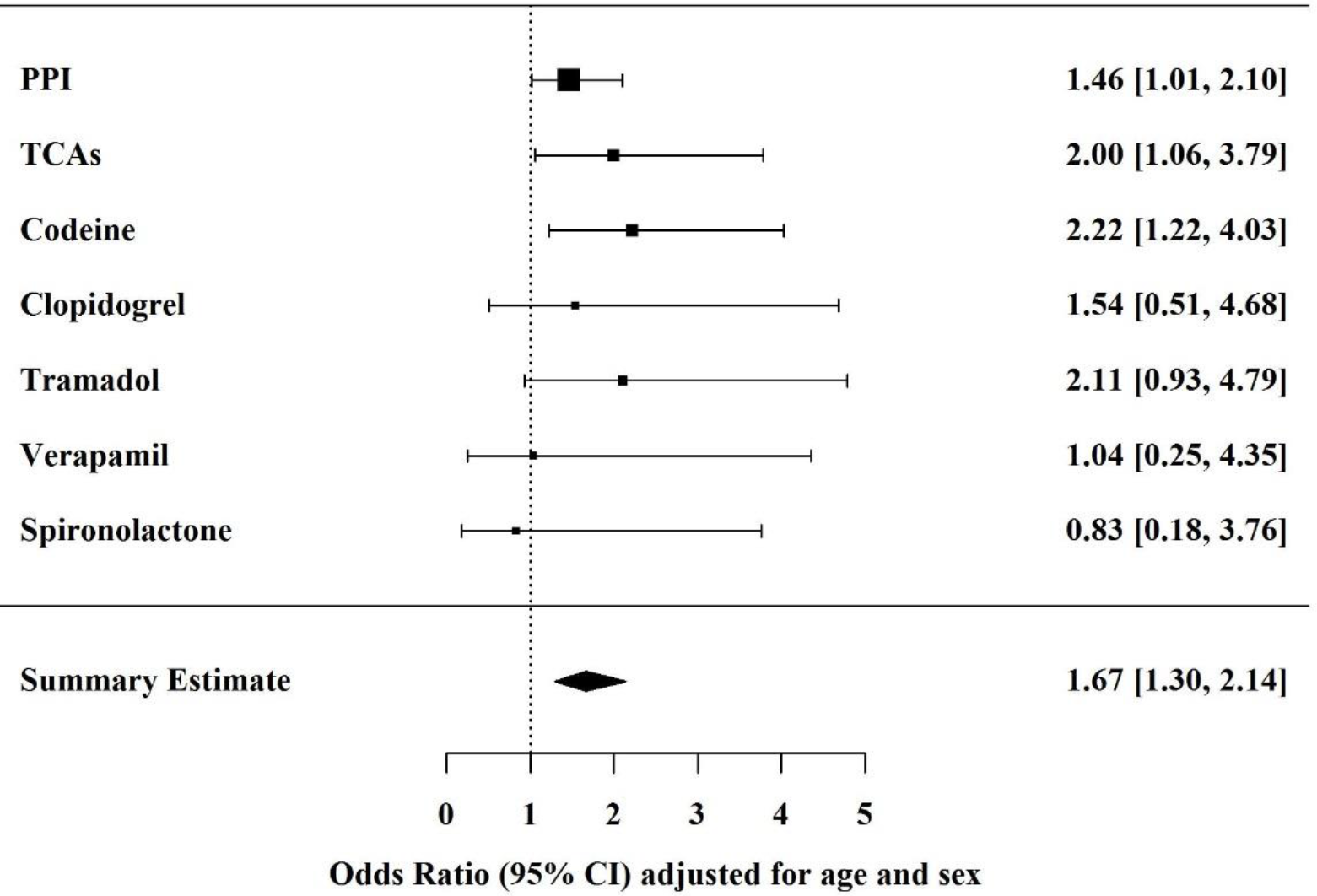
Association of individual intestinal metformin transporter inhibiting drugs with intolerance.

### Genetic variation in the gut metformin transporters and metformin intolerance

We explored the association of the intronic *SLC29A4* (rs38899348 G>A) and *SLC22A1* (M420del, R61C, G401S) SNPs with metformin intolerance. In a logistic regression model, carriers of the G allele had 1.39 [1.15-1.69, p < 0.001] times higher odds of being intolerant to metformin (unadjusted). When rs38899348 was added to a model adjusted for age, sex, weight and genetic substructure, the presence of the G allele was independently associated with metformin intolerance (OR = 1.34[1.09-1.65], p = 0.005) (Table 2). No statistically significant difference in any of the baseline phenotypes by genotype was observed (Table 3).

**Table 2.** Logistic regression model of metformin intolerance.

| Variable | OR [CI] | P |
| --- | --- | --- |
| Age at ITE (years) | 1.04[1.03-1.06] | $7.44 \times 10^{-08}$ |
| Gender | 2.31[1.72-3.10] | $2.52 \times 10^{-08}$ |
| Weight (kg) | 1.00[0.99-1.00] | 0.20 |
| Use of metformin transporter inhibiting drugs | 1.72[1.26-2.32] | $5.17 \times 10^{-04}$ |
| rs3889348 | 1.30[1.04-1.62] | 0.02 |

Gender coded as women vs men. ITE: index intolerance event

**Table 3.**
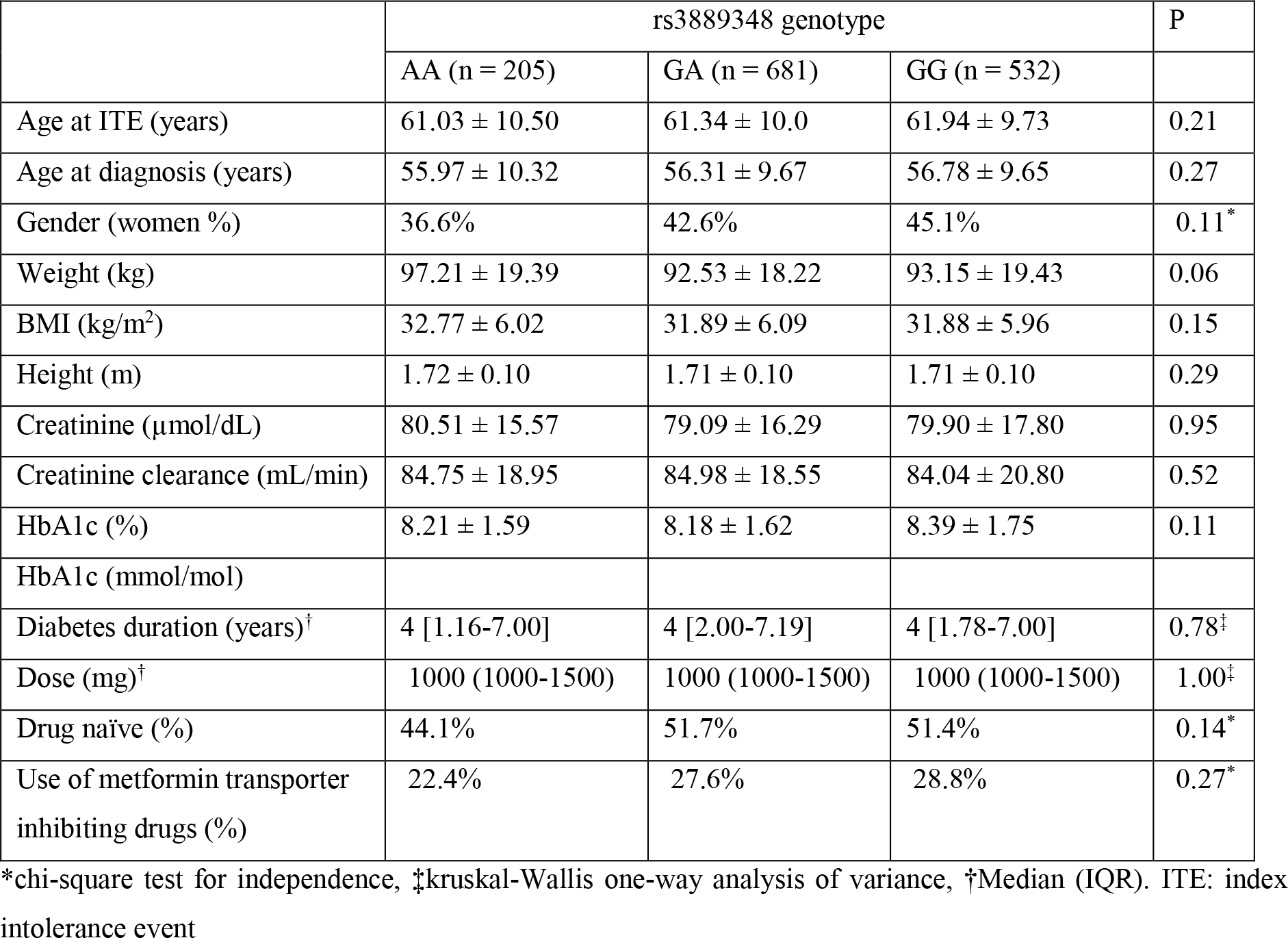
Population characteristics by rs3889348 genotype.

We then grouped subjects based on the combination of *SLC29A4* genotype and concomitant use of metformin transporter inhibiting drugs. Taking those with no risk allele and who were not treated with transporter inhibiting drugs as the reference group, carriers of one and two G alleles treated with transporter inhibiting drugs had more than two (2.44 [1.30-4.78]) and three (3.23 [1.71-6.39]) fold higher odds of intolerance, respectively, after adjusting for age, sex and weight (Table 4).

**Table 4.**
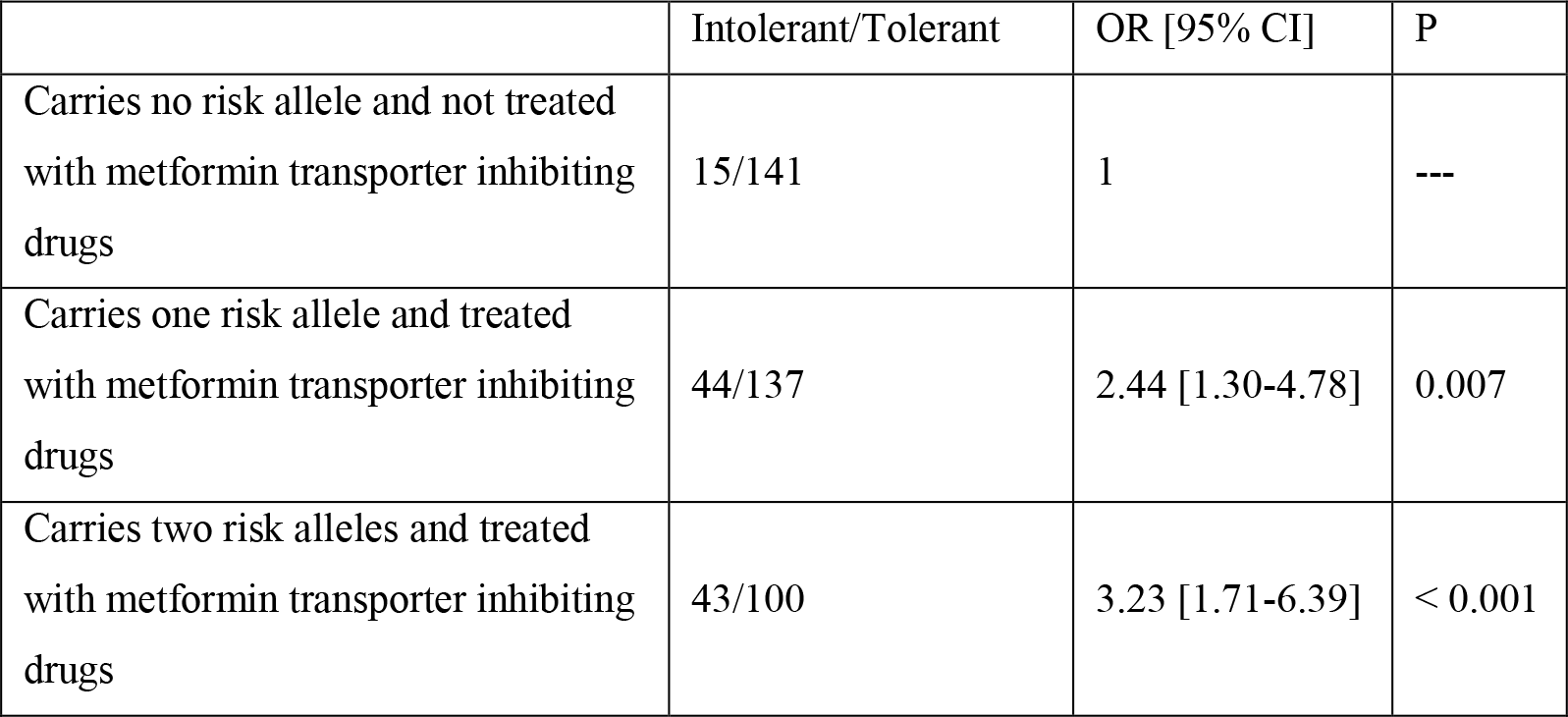
Joint effect of *SLC29A4* (PMAT) genotype and metformin transporter inhibiting drugs on metformin intolerance.

|  | Intolerant/Tolerant | OR [95% CI] | P |
| --- | --- | --- | --- |
| Carries no risk allele and not treated with metformin transporter inhibiting drugs | 15/141 | 1 | --- |
| Carries one risk allele and treated with metformin transporter inhibiting drugs | 44/137 | 2.44 [1.30-4.78] | 0.007 |
| Carries two risk alleles and treated with metformin transporter inhibiting drugs | 43/100 | 3.23 [1.71-6.39] | < 0.001 |

The association between *SLC22A1* genotypes and metformin intolerance has been previously reported (12, 32). We carried out an analysis on the association between two reduced function (R61C, G401S) and one loss of function (M420del) *SLC22A1* SNPs and metformin intolerance using a combined unweighted GRS. In a logistic regression model adjusted for age, sex, weight, genetic substructure and concomitant use of transporter inhibiting drugs, the *SLC22A1* GRS was not statistically significantly associated with metformin intolerance (OR = 1.35 [0.84-2.12], p = 0.21).

A GRS was then generated from *SLC29A4* and *SLC22A1* variants by summing up the number of risk alleles for each individual. Compared to those with no risk allele, metformin treated subjects with type 2 diabetes having two risk alleles had nearly a two-fold (1.93[1.10-3.65]) increased odds of GI intolerance. Those who carry 3 or more risk alleles had more than twice (2.15[1.20-4.12]) the odds of intolerance (Figure 2).

**Figure 2.**
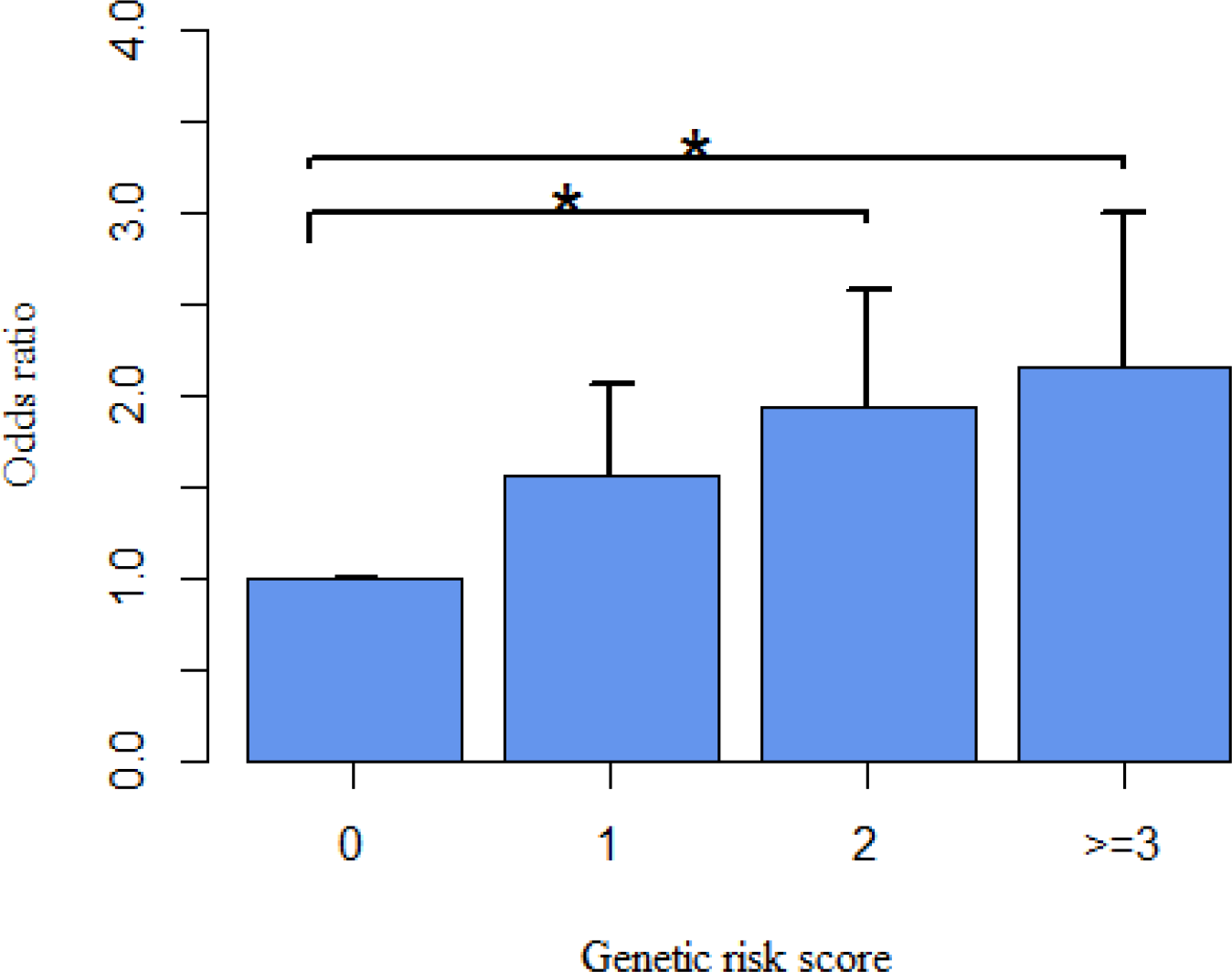
Association of a genetic risk score derived from *SLC29A4* (PMAT) and *SLC22A1* (OCT1) with metformin intolerance. OR: odds ratio; GRS: genetic risk score. Bars indicate standard errors around the mean.

### rs3889348 is associated with altered PMAT expression in the gut

Given PMAT is one of the major metformin transporters in the gut, we explored the possibility that the intronic SNP, rs3889348 is a cis-eQTL in the intestine utilizing the publicly available data set from the GTEx portal (Version V6p) (33). The G-allele of rs3889348 (associated with higher risk of intolerance) was significantly associated with lower expression of *SLC29A4* in the terminal ileum of the small intestine (β = −0.42, p = 2.1×10^−04^) and the transverse colon (β = −0.45, p = 1.4×10^-08^) (Figure 3). rs3889348 is the top cis-eQTL for *SLC29A4* in the transverse colon.

**Figure 3.**
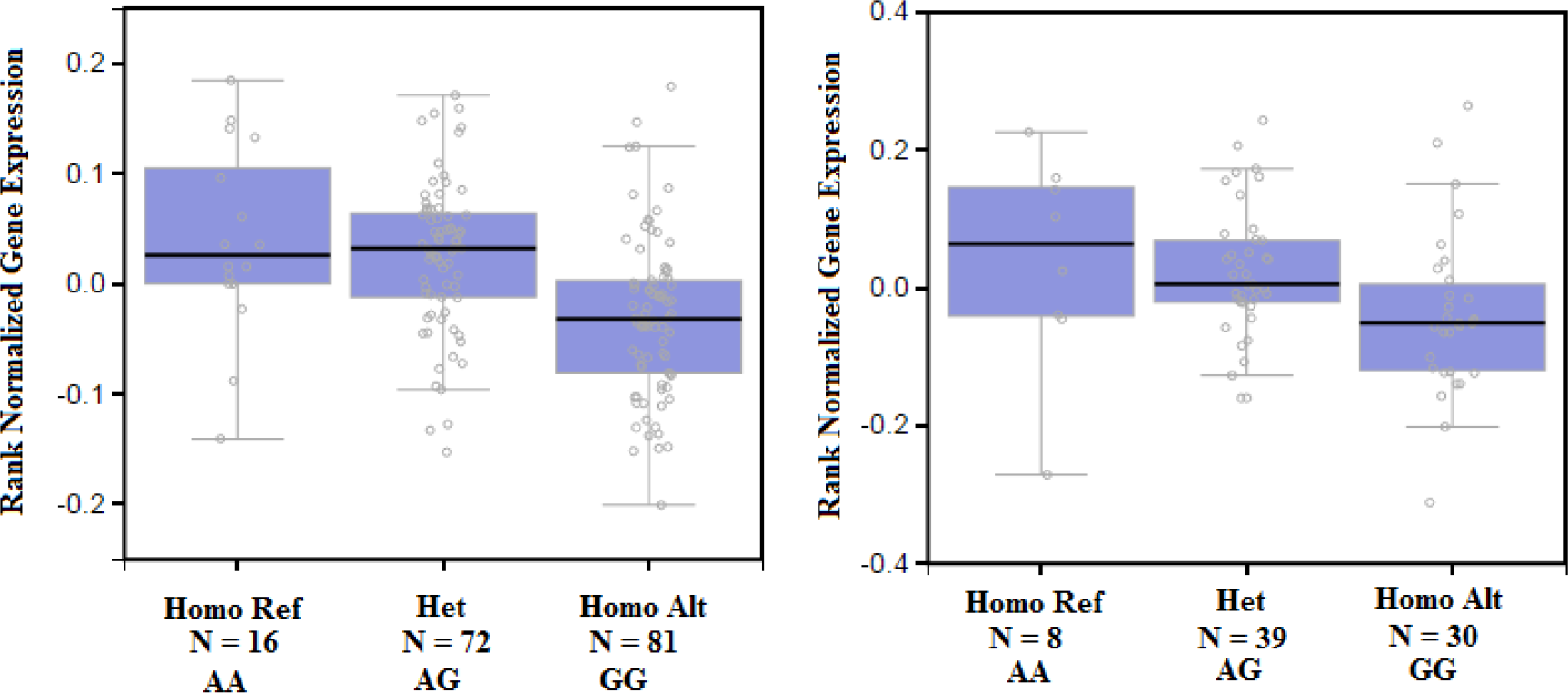
Boxplot of association between rs3889348 genotype and *SLC29A4* (PMAT) expression in the gut, colon transverse (left side) and terminal ilium of the small intestine (right side).

## Conclusions

Intestinal absorption of metformin is modulated by the function of cation transporters expressed in the gut. An association between reduced function alleles in the *SLC22A1*, encoding organic cation transporter 1, and metformin related GI side effects has been previously reported (12, 13, 34). However, the data on intestinal localization of OCT1 is ambiguous; with mixed reports suggesting in the apical (10) and basolateral (11, 36) sides. In addition to OCT1, PMAT also contributes to the intestinal absorption of metformin. PMAT is abundantly expressed in the human intestine and it is concentrated on the tips of the mucosal epithelial layer (35). Carriers of the G allele at this locus (rs3889348) had significantly reduced expression of *SLC29A4* in the gut (33). This could lead to higher luminal concentration of metformin. In this current study we demonstrated a significant association of the G allele of an intronic SNP, rs3889348, in *SLC29A4* encoding PMAT, with higher odds of GI intolerance after metformin therapy. Each copy of the G allele was associated with 1.34 times higher odds of metformin intolerance. We also show that those who carry two or more variants at either *SLC29A4* or *SLC22A1*, were two-fold more likely to have GI intolerance. Given that PMAT is apically located, this finding suggests that intolerance is driven by increased luminal concentration of metformin, rather than increased enterocyte concentration and direct toxicity to the enterocytes.

There are a number of putative mechanisms whereby increased luminal metformin may increase GI intolerance to metformin (outlined in Figure 4). Firstly, a higher concentration of metformin in the gut has been shown to inhibit uptake of histamine and serotonin leading to increased luminal concentration of these biogenic amines (13). Metformin is also shown to inhibit diamine oxidase (DAO), an enzyme that degrades histamine, at therapeutic doses (6). Biogenic amines play an important role in the GI pathophysiology. Elevated levels of serotonin and histamine in the GI tract cause GI symptoms such as nausea, vomiting and diarrhea (6, 37). Serotonin is produced mainly in the gut and stored in the enterochromaffin cells of the epithelium. Its release activates gut sensory neurons that will increase intestinal motility, secretion and sensation (37, 38). Increased colon motility and softening of stool consistency has also been observed in serotonin reuptake transporter (SERT) knock-out mice (37, 38). In addition, a recent study from the GoDARTS cohort showed association of a composite SERT genotype, 5-HTTLPR (5-hydroxy tryptamine (serotonin) transporter linked polymorphic region)/rs25531, with intolerance to metformin in subjects with type 2 diabetes (13). In this study, carriers of the low-expressing SERT S* alleles had more than 30% increased odds of metformin intolerance (OR=1.31, 95% CI 1.02-1.67, p = 0.031). Histamine is a monogenic amine stored in the enterochromaffin-like cells within the gastric glands of the stomach. Binding of histamine to the H1, H2 and H4 receptors that are highly expressed in the gut, stimulate gastric acid secretion, increase intestinal motility and smooth muscle inflammation (6).

**Figure 4.**
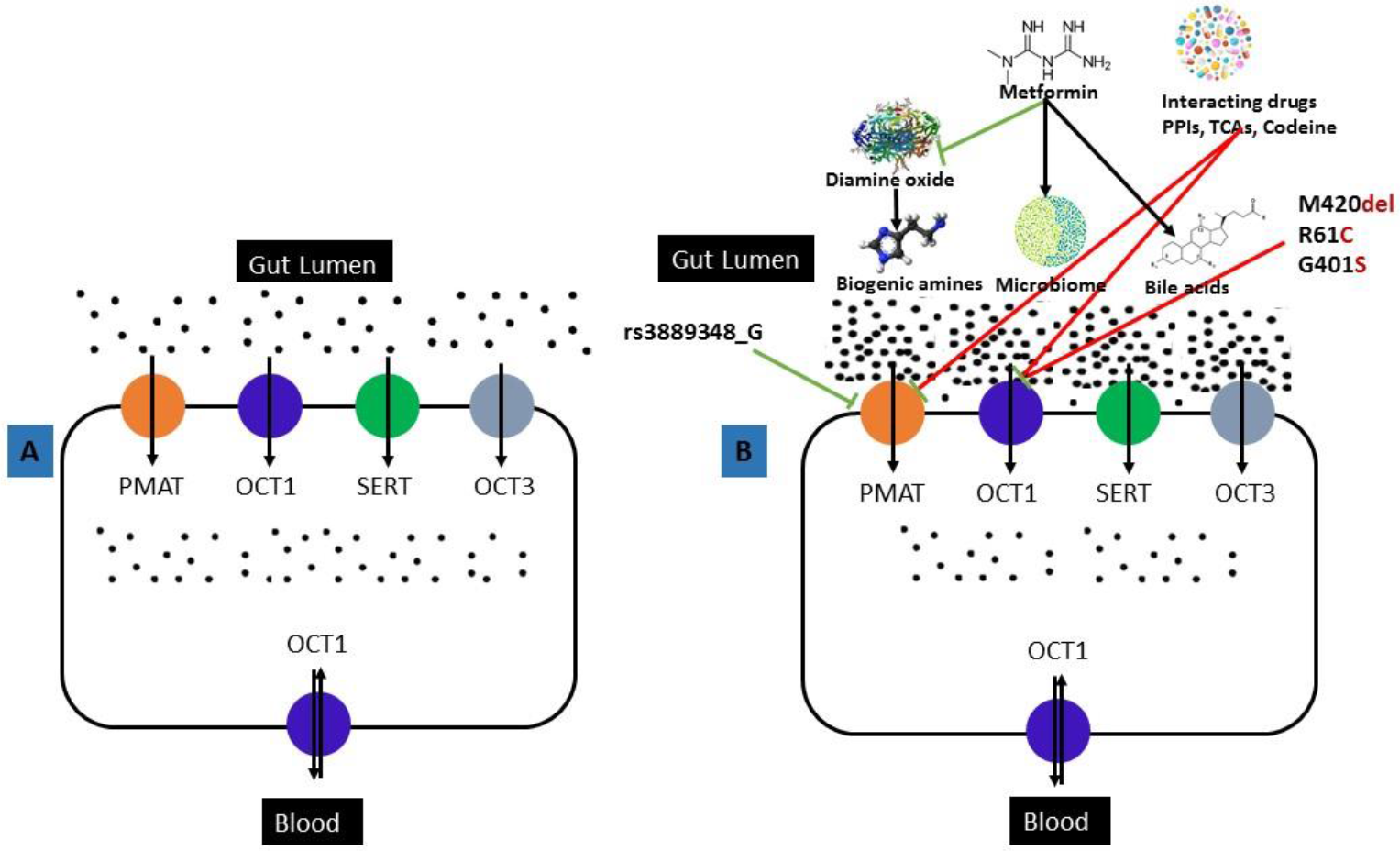
Possible mechanisms for metformin intolerance. A) Metformin is absorbed from the gut lumen via cation transporters such as PMAT, OCT1, SERT and OCT3. B) Increased level of metformin in the gut lumen is observed when metformin is taken with cation transporter inhibiting drugs such as PPIs, TCAs and Codeine. These drugs competitively inhibit metformin uptake by the cation transporters. Metformin is also shown to inhibit diamine oxide, an enzyme that metabolize biogenic amines. In addition, transport capacity of the cation transporters could be reduced in carriers of reduced function (420del, 61C, 401S in *SLC22A1*) or low expressing alleles (rs3889348_G in *SLC29A4*) and hence increased luminal metformin level. Increased level of metformin increases the level of biogenic amines, affect the gut microbiota and elevate bile acid levels. These may cause symptoms of gastrointestinal side effects.

In addition to the potential role of local concentrations of serotonin and histamine, increased luminal concentrations of metformin could also cause intolerance by other mechanisms that need to be explored. For example, intolerance could be mediated by a reduction in bile acid reabsorption in the ileum leading to elevated bile acid levels in the colon (39), which is known to cause GI disturbances (40). In addition, metformin affects composition and function of the gut microbiota favoring the growth of some species like *Akkermansia* (5, 41–43). Furthermore, increased levels of active and total GLP-1 levels in subjects with type 2 diabetes and without type 2 diabetes treated with metformin (44) have also been reported and this might increase GI side effects (45) (Figure 4).

In this study we observed increased risk of intolerance with older age, female sex, lower weight and lower creatinine levels. Concomitant use of metformin with the PPIs and TCAs also increase the risk of intolerance. These findings are largely consistent with the results of previous studies, providing further evidence for clinical practice (12, 34).

In summary, we have identified a variant that alters intestinal expression of the cation transporter PMAT (*SLC29A4*) that increases risk of metformin associated gastrointestinal intolerance, and that combined with the previously reported OCT1 variants, this genotype profile can increase odds of metformin intolerance over 2-fold. The apical location of PMAT means that reduced expression will result in increased luminal metformin concentration, suggesting that metformin intolerance is caused by this increased luminal concentration rather than increased enterocyte concentration.

## Acknowledgements

We are very grateful to all participants who took part in these studies.

